# Daylight, Daily Rhythms, and Downstream Physiology: A Translational Study in Diurnal Nile Grass Rats (*Arvicanthis niloticus*)

**DOI:** 10.64898/2026.07.06.733427

**Authors:** Antony B. Kim, Katrina Linning-Duffy, Melanie Balbach, Nolan Lucera, Marshall Delgado, Nikhil Kummur, Huishi Toh, Luisa Caldas, Lily Yan

**Author notes:** Corresponding Author: Dr. Lily Yan, Department of Psychology & Neuroscience Program 4014 Interdisciplinary Science and Technology East Lansing, MI 48824.

## Abstract

The circadian system evolved under natural light–dark cycles, while modern humans spend much of their time indoors under electric lighting that differs substantially from daylight in intensity, spectral composition, and temporal structure. How such lighting environments influence circadian system has not been systematically examined in a diurnal animal model under ecologically relevant conditions. In this study, we used the diurnal Nile grass rat (*Arvicanthis niloticus*) to assess daily locomotor rhythms across four lighting conditions designed to approximate common human exposure scenarios: rectangular daylight (D65-R; ∼5,600 lux), semi-sigmoidal daylight mimicking natural intensity dynamics (D65-S; matched peak intensity with ∼50% lower cumulative energy), fluorescent indoor light (F12; ∼150 lux), and fluorescent light supplemented with a one-hour midday daylight pulse (F12+D65-P). Using a within-subject design (n = 8), male grass rats were housed under each condition for two weeks. D65-R produced the highest daytime activity levels and the strongest day/night activity ratio, consistent with robust circadian entrainment. Despite matching peak intensity, D65-S did not yield comparable circadian outcomes, indicating that cumulative photon exposure, rather than peak intensity alone, contributes to entrainment strength. Notably, the addition of a one-hour midday daylight pulse (D65-P) partially increased circadian amplitude under otherwise fluorescent conditions, with higher periodogram amplitude relative to F12 alone. A separate cohort of males was exposed to D65-R or F12 for six weeks (n = 10/condition) to assess physiological outcomes, including metabolic and reproductive measures. Compared with the D65-R group, F12 group showed higher diabetic rate (10% *vs.* 40%) and reduced sperm mobility (45±6 vs. 19±1 %), consistent with potential downstream correlates of circadian rhythm disruption. Together, these findings demonstrate that lighting conditions characteristic of indoor environments produce weaker circadian organization than daylight-equivalent lighting in a diurnal rodent, which underscore the importance of light quality in shaping circadian rhythms and downstream physiological processes.

## Introduction

Daily light/dark cycle is the most reliable and predictable cue to entrain the circadian clocks –– an internal timing system that coordinates daily rhythms in behaviors and physiology including sleep–wake behavior, metabolic regulation, and reproductive function (Pittendrigh, 1993). This system evolved over millions of years under natural light/dark cycle, long before electric lights became available. However, in modern life, people spend ∼90% of their time indoors, resulted in a significant reduction or deficiency in daylight exposure (Klepeis et al., 2001). Natural daylight differs profoundly from typical indoor lighting in intensity, spectral composition, and temporal dynamics, all of which influence circadian entrainment and downstream physiology (Dauchy et al., 2015; Lucas et al., 2014; Roenneberg & Merrow, 2016; Wright et al., 2013).

Within the biological rhythm research community, there is growing concern that typical indoor electrical lighting provides insufficient zeitgeber strength for robust circadian entrainment (Münch et al., 2020). Adverse health effects linked to inadequate daylight exposure –– whether directly or via circadian misalignment –– range from sleep problems and impaired daytime functioning to increased prevalence of depression, obesity, diabetes, and cardiovascular disease (Burns et al., 2021; Foster and Wulff, 2005; Landvreugd et al., 2025; Maywood et al., 2006; Rijo-Ferreira and Takahashi, 2019). Wirz-Justice et al. (2021) have emphasized a gap between the importance of daylight for human well-being and the amount of controlled research on the subject, while a recent international Delphi consensus formalized 26 evidence-based public health statements on the non-visual effects of light, underscoring the need for translatable experimental evidence (Spitschan et al., 2025). Collectively, these reports call for controlled studies comparing ecologically relevant lighting paradigms –– not just bright versus dim, but outdoor-equivalent versus indoor-equivalent –– in appropriate animal models.

The paucity of animal research on the effects of natural daylight likely reflects two major challenges: light sources that faithfully mimic natural daylight, and appropriate animal models with human-like light responses. In the present study, we overcome these challenges by using a spectral light emulator to generate daylight-equivalent full-spectrum illumination; and a diurnal rodent model, the Nile grass rat (*Arvicanthis niloticus*) that responds to light in a manner similar to humans, i.e., exhibiting increased wakefulness and alertness during light exposure. The spectral light emulators and their technical specifications have been described previously (Kim et al., 2023). The effects of emulated daylight were evaluated using Nile grass rats, a well-established diurnal model that has been used in research areas including circadian rhythms and sleep, diabetes, and seasonal affective disorder (Jiang et al., 2024; Subramaniam et al., 2018; Yan et al., 2019; Yan et al., 2020). Compared to commonly used laboratory rodents that are nocturnal, i.e. active mostly at night, Nile grass rats are active mostly during daytime, therefore providing higher translational relevance for human circadian physiology and responses to lights (Jiang et al., 2024; Yan et al., 2020).

In previous studies, grass rats housed under dim light during the day developed increased depression-like behaviors, impaired spatial memory, disrupted sociosexual behavior, and had poor sleep quality compared to those housed in bright light –– paralleling symptoms of seasonal affective disorder in humans (Ikeno et al., 2016; Leach et al., 2013; Lonstein et al., 2019; Soler et al., 2018; Soler et al., 2019; Tamogami et al., 2026). Bright light therapy reversed these effects and upregulated the orexin/hypocretin system, a neuropeptide network critical for arousal and mood (Costello et al., 2023; Costello et al., 2026). Grass rats have also been used as a model for type 2 diabetes, as they can develop diet-induced diabetes when fed on a conventional laboratory rodent chow (Chaabo et al., 2010), but are protected from diabetes when maintained on a high-fiber diet (Toh et al., 2021). Unlike diet-induced obesity mouse models, which are characterized primarily by insulin resistance with only modest levels of overt hyperglycemia, diabetic grass rats can advance to complications in the heart (Schneider et al., 2018), eye (Ranaei Pirmardan et al., 2021; Toh et al., 2019), kidney (Naseri et al., 2024) and nerve (Singh et al., 2018), more closely recapitulating the natural history of human type 2 diabetes. These findings collectively support the use of Nile grass rats to investigate how indoor and outdoor lighting differentially regulate daily rhythms and physiological outcomes including metabolic regulation and reproductive function.

A key design consideration of the present study was the selection of lighting conditions as a rank-ordered set of ecologically relevant paradigms, i.e. controlled analogs of real-world human lighting environments spanning the outdoor-to-indoor continuum. This approach was intended to maximize downstream translatability, as the conditions map directly onto scenarios encountered by humans in the built environment, design programming for architects, indoor environmental quality metrics tracked by building scientists, and exposure paradigms tractable for public health assessments. Recent work by Bano-Otalora et al. (2021) in the diurnal rodent *Rhabdomys pumilio* demonstrated that bright daytime light enhances circadian amplitude in suprachiasmatic nucleus electrical activity under stable entrainment, which underscores the importance of testing graded daytime lighting conditions in diurnal species. In the present study, daily locomotor rhythms were examined under four sequential lighting conditions, and physiological outcomes were assessed in a separate cohort. The study represents an initial step toward building the comparative evidence base that the health science nexus downstream requires.

## Methods

### Animals

Nile grass rats (*Arvicanthis niloticus*) were obtained from a breeding colony at Michigan State University (Yan et al., 2020). Animals in the colony were kept in a 14:10 hour light/dark (LD) cycle with lights-on at 0600 h and lights-off at 2000 h, defined as Zeitgeber time (ZT) 0 and 14, respectively. Light was supplied by ceiling lights (GE current, 14BDT8/G4/840, 4000 k); illuminance was around 300 lux in the center of the room. Food (Prolab 2000 #5P06, PMI Nutrition LLC, MO, USA) and water were available *ad libitum.* A metal hut was provided in each cage for shelter and enrichment. All experimental procedures followed the guidelines of the NIH Guide for the Care and Use of Laboratory Animals (NIH Publication No. 80-23) and were approved by the Institutional Animal Care and Use Committee of Michigan State University.

### Daytime lighting conditions in the study

The experimental animals were housed in different daytime light conditions (Fig. 1), provided by a spectral light emulator (Telelumen Inc., Moorpark, CA, USA) installed above each cage. The custom 8-channel spectral light emulators are equipped with 6 narrowband and 2 broadband LEDs that cover the wavelength range of 360-658 nm to provide appropriate spectral fidelity to stimulate all known rodent retinal photopigments, including UV-type S cones. Seven channels were specified for photopigment stimulation, and one red channel, outside the rodent photopigment response range, was included to facilitate subjective-night experimental procedures. The emulators incorporate pulse width dimming (up to 1000:1) and a frame-rate control (up to 1 Hz), enabling precise modulation of intensity and temporal dynamics (Kim et al., 2023). The emulators were operated at approximately 85% of maximum rated output to preserve LED longevity over the multi-month experimental period.

**Figure 1.**
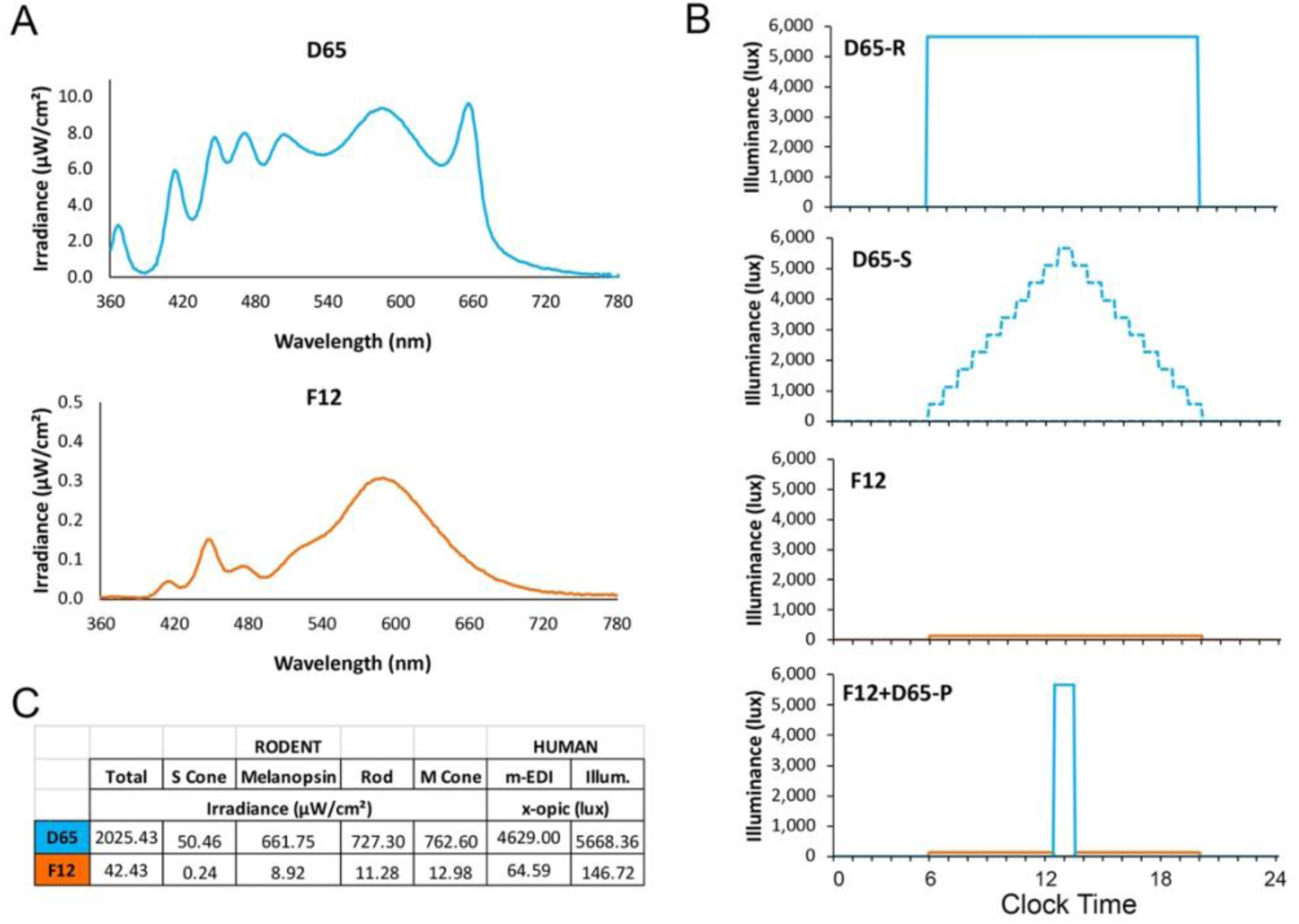
Lighting conditions in the study. A, spectral irradiance measurements for the D65 and F12 conditions. B, the 14:10 h light/dark paradigms with 4 different daytime light conditions: rectangular D65 (D65-R), a semi sigmoidal D65 (D65-S), F12, and F12 with one-hour D65 pulse at midday (F12+D65-P). C, irradiance of D65 and F12 was calculated in total and by individual photoreceptor type. Melanopic EDI (m-EDI) and photopic illuminance in the right columns are human-centric measures and were included for reference purposes only.

Two spectra were used: International Commission on Illumination CIE standard illuminant daylight 6500 K (D65) and CIE standard illuminant fluorescent #12 (F12) to simulate typical outdoor and indoor light conditions, respectively (Fig. 1A). D65 is the CIE standard illuminant for average natural daylight, a broad reference for the spectral composition of natural daylight. The intensity of D65 used in the present study (∼5,600 lux) matches the operative definition of “overcast daylight”, which is heavily diffuse sunlight filtered through standard cloud cover, ranging from about 1,000 to 10,000 lux (Egan and Olgyay, 2001; Reinhart, 2020; Wirz-Justice et al., 2021). The emulated daylight of D65 at ∼5,600 lux is hereafter referred as “daylight”.

Four housing conditions (Fig. 1B), all set to 14:10 LD cycle, were tested: (1) D65-R (rectangular daylight): D65 at ∼5,600 lux intensity throughout the day, serving as the control condition and representing sustained outdoor overcast daylight condition. (2) D65-S (semi-sigmoid daylight): D65 gradually reaching peak intensity of ∼5,600 lux at midday then dimming to complete darkness by the end of the day, modeling the natural dynamic of daylight brightness. Peak intensity was matched to D65-R, but the sinusoidal envelope delivered approximately 50% less total energy over the subjective day. (3) F12 (fluorescent indoor): F12 at ∼150 lux throughout the day, representing the standard illuminance design target for modern office spaces dominated by electrical lighting. (4) F12+D65-P (fluorescent with midday daylight pulse): the same F12 condition with a 1-hour D65 pulse (∼5,600 lux) at midday, simulating the experience of an office occupant who steps outside for a one-hour walk during lunch.

Light was measured using a calibrated spectroradiometer (MSC15, Gigahertz-Optik) at bottom-of-cage level. The photic input for each rodent photoreceptor type was calculated using the Rodent Toolbox v.2 (Lucas et al., 2023), and corresponding human-centric photopic illuminance and melanopic EDI (mEDI) lux values were calculated using the CIE S 026 α-opic Toolbox (CIE, 2020), as described previously (Kim et al., 2023), and reported in Fig. 1C.

### Experiment 1: Impact of daytime light conditions on daily locomotor activity

Male grass rats (3-4 months old, n = 8) were singly housed under four lighting conditions described above sequentially. Following the first 5 days of habituation period, the animals were housed in each condition for two weeks. In-cage locomotor activity was monitored continuously using an Actimeter system, and an Internet of Things sensor platform (Geocene Inc., Berkeley, CA) as in previous studies (Kim et al., 2023). Locomotor activity data were processed using ClockLab (Actimetrics, Inc. IL, MSU) to generate actograms and for quantitative analysis. Higher level of activity (∼30-40% increase) was observed on each Thursday and Friday likely due to the weekly cage change on Thursday in late afternoon, therefore those two days in each week were excluded for the analysis of daily activity and rhythms. For the 2-week period under each condition, 10 days of data were used to analyze daily activity, amplitude of periodograms, day/night activity ratios, and entrainment precision (standard deviation of as described in the day-to-day variability of activity onset and offset) as in previous studies (Leach et al., 2013). A single-component (24-h) cosinor model was also fit to each animal’s daily activity by ordinary least squares to derive the MESOR, amplitude, and acrophase (Nelson et al., 1979; Cornelissen, 2014). Rhythm regularity and fragmentation were further characterized by inter-daily stability and intra-daily variability (Van Someren et al., 1999).

### Experiment 2: Impact of daytime light conditions on physiological outcomes relevant to metabolic regulation and reproductive function

A separate cohort of male grass rats (3-4 months old) were housed in either D65-R or F12 as in experiment 1 (n = 10/condition) for 6 weeks, followed by the assessment of physiological outcomes including body weight, blood glucose level, weight of perigonadal fat pad/white adipose tissue, weight of testis and epididymis, sperm concentration and mobility. To ensure all the measurements were taken within the same time window 0900-1000 hr (ZT3-ZT4), samples were collected in two days. Animals were euthanized with CO_2_ at 20% of the chamber volume per minute for 4 to 5 minutes until cessation of breathing, which was consistent across animals, immediately followed by the measuring of body weight, and blood glucose measures from right atrium using a glucometer (MicroTech, GoCheck2). Perigonadal fat pad, testes, seminal vesicles, and epididymis were dissected and weighed. Epididymis collected from each animal were kept on a 40°C heating pad while transported to another building on campus for sperm analysis.

Epididymis collected from the first day were excluded due to technical reason, and only the tissue/data from the second set (n = 5/condition) were used for the sperm analysis.

### Sperm analysis

Sperm were isolated by incision of the cauda epididymis in 1 ml mKRB medium (in mM: 94.6 NaCl, 4.8 KCl, 1.7 CaCl_2_, 1.2 KH_2_PO_4_, 1.2 MgSO_4_, 20 HEPES, 5.6 glucose, 0.5 sodium pyruvate, pH 7.4 adjusted at 37°C with NaOH) prewarmed at 37°C. After 15 min swim-out at 37°C, sperm were diluted 1:50 in mKRB medium, 25 µl sperm suspension were loaded on a 100 μM Leja slide (Hamilton Thorne) and placed on a microscope stage at 37 °C. Sperm motility was assessed using computer-assisted sperm analysis (CASA) via a Hamilton–Thorne digital image analyzer (IVOS II, Hamilton Thorne Research, Beverly, MA) with the following parameters: 30 frames, frame rate: 60 Hz, cell size: 30–170 μm^2^. Sperm movements of 10 fields of at least 1000 sperm were examined. Sperm were considered motile when presenting straightness (STR) ≥ 80 % and average path velocity (VAP) ≥ 5 μm/s and considered progressively motile when presenting STR ≥ 80 % and VAP ≥ 50 μm/s.

### Quantification and Statistical Analysis

Locomotor activity and rhythm parameters were computed in Python using custom scripts built on the NumPy, SciPy, and pandas libraries. Statistical analysis performed using the pingouin and statsmodels packages. The effects of lighting condition on locomotor activity and rhythm parameters were evaluated by one-way and two-way repeated-measures ANOVA, with light condition and, where applicable, time or phase (day vs. night) as within-subject factors. These analyses were applied to the hourly activity profiles, total day and night activity, periodogram amplitude, the day/night activity ratio, MESOR, amplitude and acrophase in Cosinor fitting, entrainment precision, inter-daily stability and intra-daily variability. Significant effects of lighting condition were followed by Holm-corrected post hoc pairwise comparisons. For the physiological outcomes, unpaired t-tests were used to compare body weight, perigonadal white adipose tissue, blood glucose, sperm concentration and motility between the D65-R and F12 lighting conditions. Statistical significance was set at p < 0.05.

## Results

### Impact of daytime light conditions on daily locomotor activity

A clear diurnal pattern was observed in all 8 animals, with more activity during light phase than in dark phase (Fig. 2). Comparing the four different housing conditions, the overall activities appeared to be higher in D65 conditions than in the F12 conditions in most of the animals. In the last condition of F12, a distinct bout of higher activity during the 1hr D65 exposure around noon was observed in all 8 animals. The impact of light conditions on daily locomotor activity was confirmed by quantitative analysis (Fig. 3). Comparing the hourly activity profile under the 4 lighting conditions (Fig. 3A), a two-way repeated measure ANOVA revealed a significant effect of light condition (F_3,21_= 4.7, p < 0.05), a significant effect of time (F_23,161_ = 22.17, p < 0.01), and a significant interaction between the two factors (F_69,483_ = 5.26, p < 0.05). One-way ANOVA revealed significant effect of light condition in most hours during the day (at 7 and 17, p < 0.05; from 8-16, p < 0.01), but not in any of the night hours (p > 0.05). Total activities during day and night in the 4 conditions showed a similar pattern (Fig. 3B). A two-way repeated measure ANOVA revealed a significant effect of phase (day *vs.* night, F_1,7_ = 35.32, p < 0.001), a significant effect of light condition (F_3,21_ = 5.09, p < 0.01), and a significant interaction between the two factors (F_3,21_ = 6.59, p < 0.01). A repeated measure ANOVA showed a significant effect of lighting condition only during the day (F_3,21_ = 9.36, p < 0.001), but not at night (F_3,21_ = 0.38, p = 0.76). Post-hoc pairwise comparison of activities during the day revealed higher activity in D65-R, than in the other 3 conditions (p < 0.05 in all 3 cases). Amplitude of periodogram and day/night activity ratio were used to assess the robustness of the daily rhythms. Light condition had a significant effect on the amplitude of periodogram (Fig. 3C, repeated measure ANOVA, F_3,21_ = 3.09, p < 0.05), with the amplitude in F12+D65-P higher than that in D65-S (p < 0.01). Lighting condition also had a significant effect on day/night activity ratio (Fig. 3D, F_3,21_ = 3.08, p < 0.05), with a significant difference between D65-R and F12 (p < 0.01) and a marginally significant difference between F12 and F12+D65-P (p = 0.08). A Cosinor fitting analysis revealed a significant effect of light conditions on the MESOR (Fig. 3E, F_3,21_ = 4.67, p < 0.05), amplitude (Fig. 3F, F_3,21_ = 6.65, p = 0.01), and acrophase (Fig. 3G, F_3,21_ = 2.3, p < 0.05) of the daily rhythm. The amplitude in F12 was significantly lower than that in D65-R (Fig. 3F, p < 0.05), with acrophase significantly later than the two D65 conditions (Fig. 3G, p < 0.05). There was no significant effect of light condition on the precision of activity onset (Fig. 3H) or offset (data not shown). Inter-daily stability in F12+D65-P condition was significantly higher than in D65-S (Fig. 3I, p < 0.05), and marginally higher than that in F12 (p = 0.06). Intra-daily variability appeared to be lower in the D65 conditions than that in the F12 conditions (Fig. 3J), but the effect didn’t reach statistical significance.

**Figure 2.**
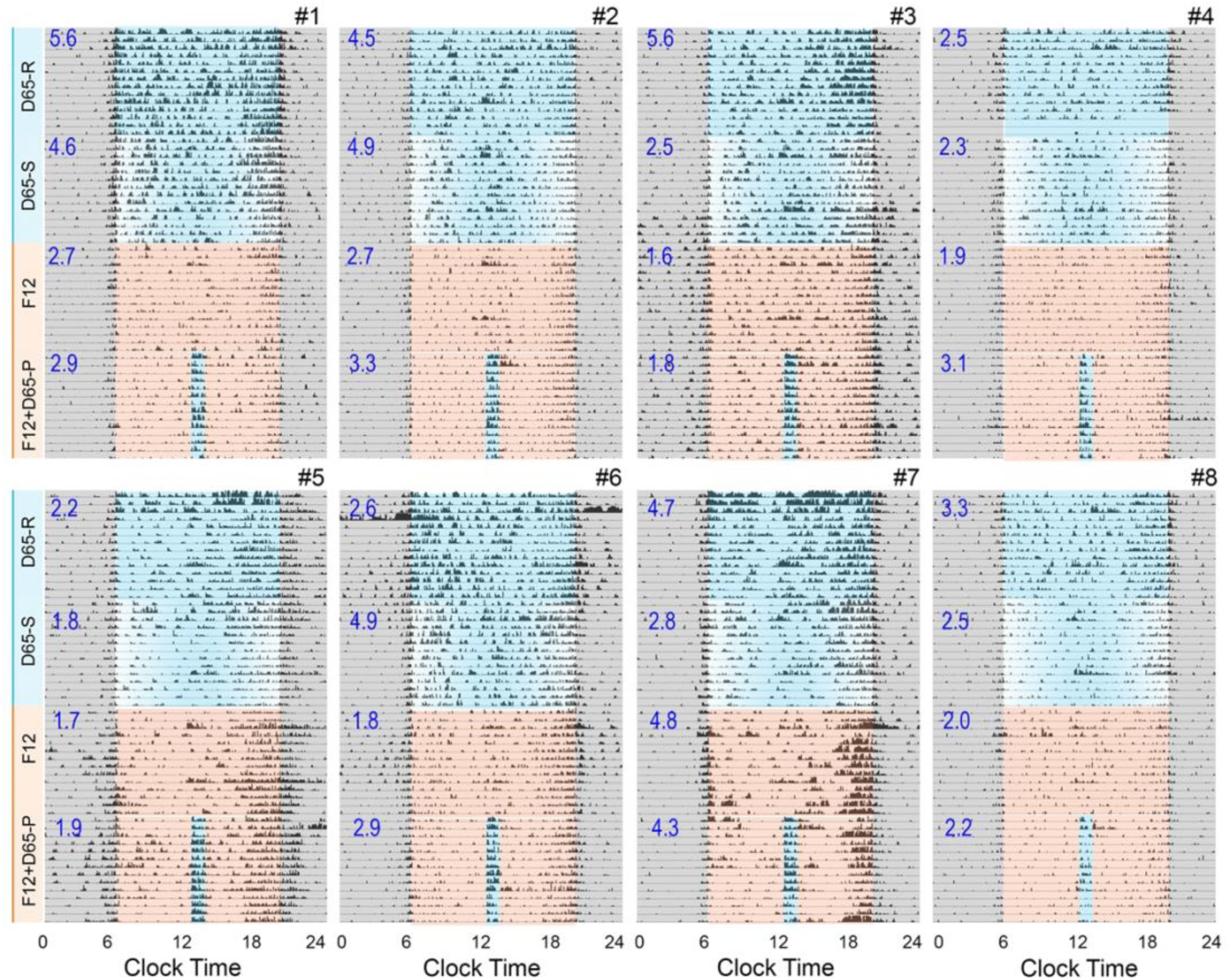
Actograms of all 8 animals showing daily rhythms of locomotor activities under each light condition. Animals were first entrained under D65-R, next D65-S, then F12, and lastly F12+D65-P. Numbers shown on the actograms represent the day/night activity ratio under each light conditions.

**Figure 3.**
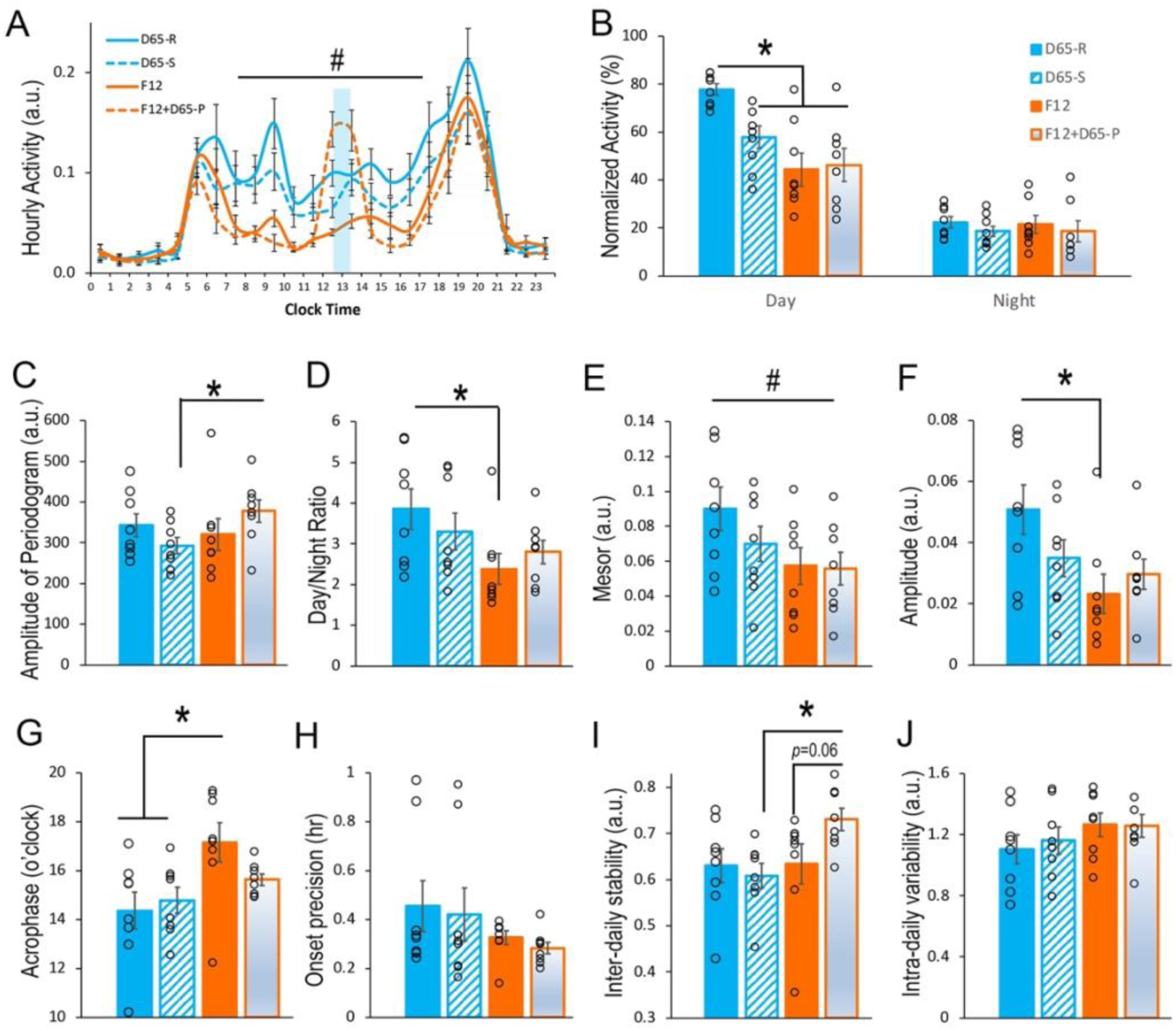
Quantitative analyses of daily activity under each condition including daily activity in hourly bins. (A), total activity in day and night (B), amplitude of periodogram (C), day/night activity ratio (D), MESOR (E), amplitude (F) and acrophase (G) of a cosinor analysis, onset precision (H); I, inter-daily stability; and J, intra-daily variability. Data are shown as mean ± SEM (n = 8). Open circles in bar graphs represents individual data points.^#^ indicates significant effect of condition in repeated measure ANOVA, p < 0.05; * indicates significant difference in post-hoc pairwise comparison, p < 0.05.

### Impact of daytime light conditions on metabolic outcomes

There were no significant differences in body weight (Fig. 4A), perigonadal white adipose tissue (Fig. 4B), or blood glucose level (Fig. 4C) between the two groups housed in either D65 or F12 condition for 6 weeks. Blood glucose level showed a clear dichotomy (Fig. 4C). Although most animals had glucose levels below 150 mg/dL, 5 out of 20 animals had glucose measurements in the 400–500 mg/dL range, suggestive of a diabetic state (Subramaniam et al., 2018). 4 out of the 5 diabetic animals are from the F12 housing condition.

**Figure 4.**
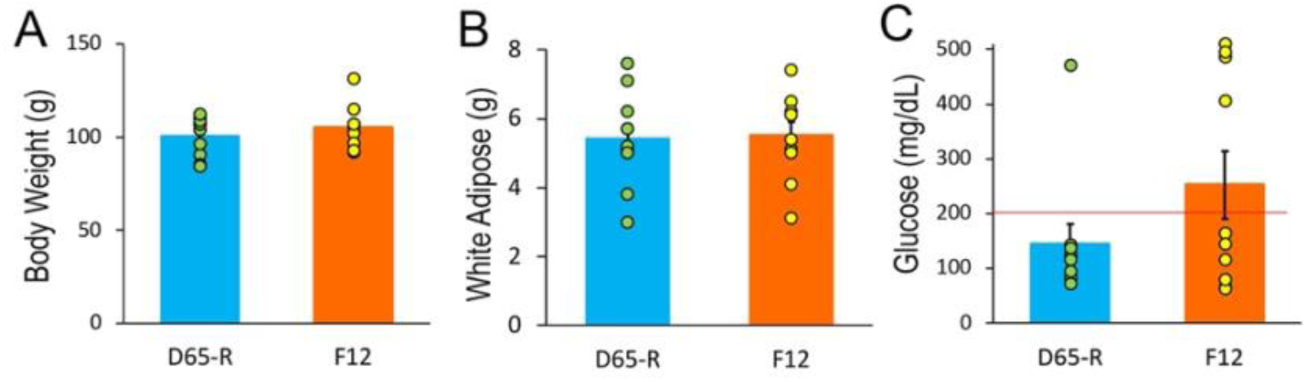
Body weight (A), perigonadal white adipose tissue (B) and non-fasting blood glucose level (C) in animals housed under D65-R or F12 lighting conditions for 6 weeks. Data are presented as mean ± SEM (n = 10), with individual animals shown as filled circles. Red horizontal line in panel C indicates a blood glucose level of 200 mg/dL.

### Impact of daytime light conditions sperm production and motility

Sperm counts between the two groups were not different between the two conditions (Fig. 5A), suggesting that spermatogenesis is unaffected by the lighting conditions. However, both the percent of motile (Fig. 5B, p < 0.01) and progressively motile sperm (Fig. 5C, p < 0.05) was significantly lower from animals in the F12 housing conditions than the D65-R housing conditions, revealing a so far undetected effect of lighting on sperm function.

**Figure 5.**
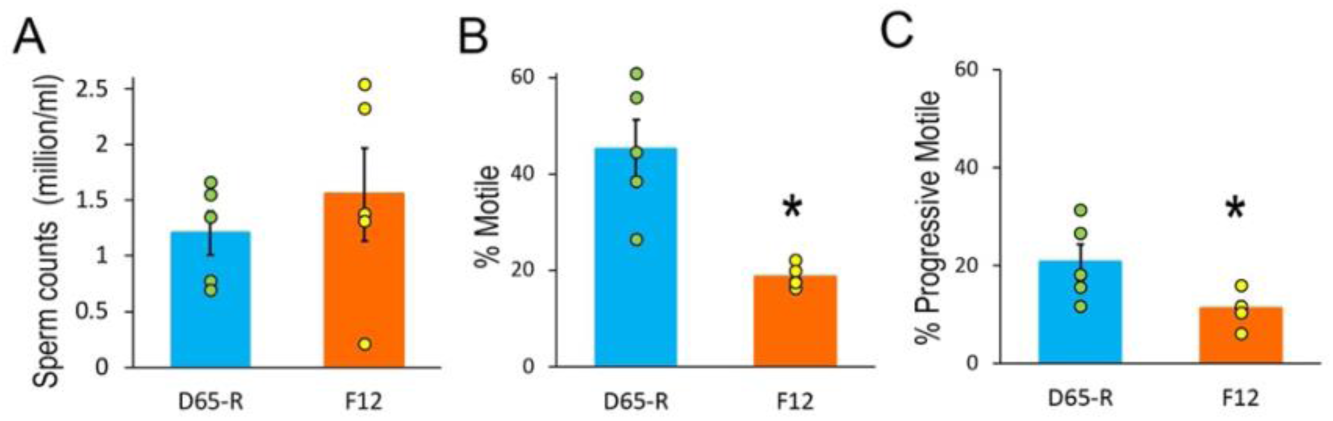
Sperm counts (A), % motile (B) and % progressive motile (C) of animals housed in D65-R or F12 for 6 weeks. Data are shown as mean ± SEM (n = 5), with individual data points overlaid and shown as filled circles. * indicates p < 0.05.

## Discussion

The present study examined the effects of ecologically relevant lighting conditions on daily locomotor rhythms and physiological outcomes in a diurnal rodent model. The locomotor activity results revealed that: (1) emulated daylight (D65) produced more robust diurnal activity patterns than fluorescent light (F12) at typical indoor illuminance; (2) a sinusoidal D65 intensity profile (D65-S), despite matching D65-R in peak intensity, did not produce equivalent rhythm outcomes; and (3) a single one-hour midday D65 pulse superimposed on F12 (F12+D65-P) partially rescued daily rhythm amplitude (Fig. 2 & 3). Physiological outcomes were compared between animals housed in either D65-R or F12, which showed that diabetic (non-fasting blood glucose > 200 mg/dL) animals were asymmetrically concentrated in the F12 than in D65 condition, 40% vs. 10% (Fig. 4); and sperm motility was significantly reduced in the F12 than in D65-R (Fig. 5).

### Lighting Conditions as a Rank-Ordered Translational Framework

A deliberate design feature of this study was the selection of lighting conditions as a graded, rank-ordered comparison across the outdoor-to-indoor continuum: sustained outdoor light condition (D65-R), outdoor light with naturalistic temporal dynamics (D65-S), a standard electrically lit office (F12), and an office worker’s day punctuated by a midday outdoor walk (F12+D65-P). By anchoring conditions to CIE standard illuminants (D65 and F12), internationally recognized spectral reference points, the present results can be contextualized across laboratories and translated into the photometric terms used in the lighting and building industries. This is consistent with the use-inspired basic research framework (Stokes, 1997), in which fundamental questions about circadian photoreception are pursued with attention to the translational pathway.

It is worth contextualizing the intensity used here against the full range of sunlight. The ∼5,600 lux delivered in the D65 conditions sits at the range of overcast daylight, well below the 100,000–120,000 lux of direct sun. Even at this overcast daylight level, it produced significantly higher diurnal activity and rhythm amplitude than fluorescent light (F12) at 150 lux. The effects reported here therefore likely understate what sunlight would produce, and the biological gap between typical indoor and outdoor lighting is, in practice, larger than these data capture.

The relatively lower intensity in D65 likely also contributed to individual differences in the light responses that sunlight would provide. Although the majority (6 out of 8) showed higher level of activity in D65-R than in F12, 2 of them (#5, #7 in Fig. 2, Suppl. Fig. 1) had comparable level of activities between the two conditions. It is noteworthy that in the present study, all animals progressed through light conditions in a fixed order without counterbalancing. Therefore, we cannot rule out the possibility that the reduction in activity from D65-R to later conditions in some of the animals reflect age-related decline or other unmeasured environmental influence. However, in a previous study in which grass rats were initially housed in F12 and then switch to D65-R, daytime activity was also found higher in D65-R than in F12 (Kim et al., 2023). Together, these findings are consistent with the interpretation that grass rats exhibit higher activity level under daylight-equivalent lighting than under indoor fluorescent light.

### D65-R Versus D65-S: The Role of Cumulative Photon Dose

Following the D65-R condition, D65-S condition was designed to mimic the natural temporal envelope of daylight, with gradual dawn, peak noon intensity, and gradual dusk. The circadian system evolved under precisely this kind of graded intensity profile, and researchers have hypothesized that twilight transitions serve as chronobiological signals in their own right (Danilenko et al., 2000; Roenneberg and Foster, 1997). In Syrian hamsters, simulated twilight transitions expanse the range of entrainable periods and enhance the precision of activity timing relative to rectangular cycles delivering equivalent total photon dose (Boulos et al., 1996; Boulos and Macchi, 2005), This effect likely reflects the temporal structure (Usui, 2000) and spectral dimension (Spitschan et al., 2017; Walmsley et al., 2015) inherent to twilight, which provides circadian timing cues independent of total energy or brightness.

In the present study, the D65-S condition did not outperform D65-R on any measured parameter of rhythm robustness. It’s noteworthy that D65-S delivered approximately 50% less cumulative radiant energy than D65-R over the subjective day. The data therefore suggest that cumulative photon dose may matter as much as, or more than, the temporal shape of the intensity profile. In other words, the circadian system may integrate light exposure over time in a manner that gives substantial weight to total energy, not just peak intensity. This aligns with independent electroencephalographic evidence in the same species, where daytime wakefulness rose stepwise across 150, 300, and 1,000 lux and daytime sleep fell accordingly (Tamogami et al., 2026). If more light buys more wakefulness in a graded fashion, then a profile that delivers half the total energy, as D65-S did, would be expected to come up short, which is what was observed.

Another important consideration is that the D65-S condition reproduced the intensity trajectory of twilight but not its spectral (chromatic) changes, unlike earlier studies using natural outdoor daylight (Daan and Aschoff, 1975). Given evidence that the circadian system is sensitive to the chromatic signature of twilight (Mouland et al., 2019; Walmsley et al., 2015), a fixed spectrum may not capture the full stimulus underlying twilight’s reported potency. The literature on twilight is derived largely from nocturnal models, and it remains unclear whether twilight sensitivity and dose integration operates similarly in diurnal mammals. Caution is warrant, as comparative studies indicate that twilight does not function as a uniformly stronger zeitgeber across species. For example, nocturnal mice and hamsters respond to matched twilight conditions in opposite directions (Comas and Hut, 2009). Mechanisms characterized in nocturnal rodents may therefore not generalized to diurnal species such as grass rats. Elucidating how circadian system responds to twilight has practical significance for building design, as it speaks to whether architects should prioritize sustained high-intensity daylight, temporally graded exposure mimicking dawn and dusk, full spectral emulation of twilight, or some combination of the three.

### F12 and the Indoor Light Deficit

Under the F12 condition, diurnal activity levels and the day/night activity ratio (Fig. 3D) were significantly reduced relative to D65-R, indicating a weaker distinction between active and rest phases. Cosinor analysis further revealed a lower amplitude in F12 than in D65-R (Fig. 3F), and a later acrophase in F12 than in both D65 conditions (Fig. 3G), suggesting reduced rhythm robustness and a modest phase-delayed under F12. These results are consistent with a growing body of field dosimetry data quantifying the gap between indoor lighting and circadian photostimulation thresholds. The Brown et al. (2022) consensus recommended a minimum daytime exposure of 250 melanopic EDI. However, wearable dosimetry studies report that the median corneal-plane melanopic EDI in office settings is only 51 lux (Stefani et al., 2024), with the mean inflated to ∼300 only by brief outdoor spikes, and no office building studied to date has met the 250 mEDI threshold for the full working day (de Vries et al., 2025). What the dosimetry data suggests by association, the present study demonstrates by experimental manipulation in a diurnal mammalian model: standard indoor lighting produces a measurably weaker circadian signal than outdoor lighting. Additionally, our previous work has demonstrated that spectral composition matters independently of intensity: a daylight-equivalent spectrum (D65) produced tighter circadian entrainment than a fluorescent-equivalent spectrum (F12) at matched melanopic irradiance, and direct S-cone stimulation produced the greatest and longest-lasting wakefulness-promoting effects (Kim et al., 2023). An important spectral constraint warrants discussion is that the F12 provides virtually no stimulation of the UV-type S-cone in rodents but stimulates S-cone pathways in humans, representing a potential limitation for direct translational interpretation.

### The Midday Daylight Pulse: A Partial Rescue

Following the midday light pulse, a distinct activity bout during the exposure was visible in all 8 animals (Fig. 2), consistent with our previous finding of brighter light-induced increase of daytime activity in which the highest increase in activity occurred in the first hour, then gradually decline toward the baseline in the 3^rd^ hour of light exposure (Kim et al., 2023). In the present study, a single one-hour midday exposure of daylight (D65-P) was sufficient to partially restore the amplitude of daily rhythms. The day/night activity ratio (Fig. 3C), along with the amplitude (Fig. 3F) and acrophase (Fig. 3G) of the cosinor fit, reached levels comparable to those observed in D65 conditions. Beyond increasing activity level (positive masking), D65-P also improved synchronization of the daily activity rhythm, as indicated by a tighter acrophase clustering (Fig. 3G) and greater inter-daily stability (Fig. 3I). Besides the rhythm outcomes, a daily one-hour morning walk outdoors or just spending more time outdoors could produce greater improvement in depression self-ratings than artificial light, indicating that leaving the building itself is therapeutic (Rosenthal et al., 1984; Wirz-Justice et al., 1996). The translational implication is straightforward: for a worker housed under standard fluorescent lighting, even a brief midday outdoor exposure may meaningfully improve circadian health. This carries implications for workplace design, as well as for public health messaging. The present study provides controlled animal model evidence supporting what has been, until now, primarily an observational claim.

### Physiological outcomes

Beyond circadian and mental health outcomes, natural daylight has been shown to improve glucose control in individuals with type 2 diabetes (Harmsen et al., 2026) and increase the success rate of in vitro fertilization (Vandekerckhove et al., 2016). When grass rats were first brought to laboratory, many became diabetic. Dietary change obviously could be the main driving force, but the potential contribution of switching from daylight in their natural habitat to fluorescent light in laboratory has never been examined. The case for light as the culprit is not without precedent: in another diurnal rodent, the sand rat, animals housed indoors under artificial light developed type 2 diabetes while those kept outdoors under natural conditions on the same diet did not (Bilu et al., 2019). Nile grass rats have been established as a model of type 2 diabetes (Subramaniam et al., 2018). In this model, non-fasting random glucose greater than 200 mg/dL is generally considered as diabetic. A recent study has found that non-fasting, but not fasting, blood glucose correlates with the oral glucose tolerance test outcome, supporting non-fasting glucose as a more informative indicator of glucose metabolism and insulin sensitivity (Anderson et al., 2023). The same study found no significant hour-to-hour variation in non-fasting glucose across a 12-hour interval from morning to early evening. Accordingly, in the present study, blood glucose was measured in non-fasting animals around midday. It’s noteworthy that blood was collected following CO2 euthanasia, which may influence blood glucose level. In a follow-up experiment, glucose was measured from tail vein under isoflurane anesthesia in grass rats housed under D65 or F12 conditions for 4 weeks. Diabetes (glucose > 200mg/dL) was detected in 1 of 7 animals in D65-R and 4 out of 8 in F12 (Suppl. Fig. 2). Together, these findings support daylight deficiency as a risk factor for diabetes (Fig. 4; Suppl. Fig. 2). Indeed, a recent study examined a cohort of type 2 diabetes patients exposed at daylight through windows vs. constant artificial lighting during office hours (Harmsen et al., 2026). It was found that daylight exposure had a positive metabolic impact, keeping blood glucose in the normal range and promoting greater reliance on fat oxidation.

Studies on the effect of light conditions on sperm function have been focused mostly on photoperiod, in which long photoperiod (16 hrs) has been shown to improve sperm motility in animals with or without strong seasonal breeding pattern, e.g., ram (Zhu et al., 2024), or Sprague-Dawley rats (Olayaki et al., 2008). Although effects of daylight have not been directly studied in humans, exposure to brighter daylight has been shown to increase the success rate of IVF (Vandekerckhove et al., 2016). By analyzing nearly ten thousand cycles of IVF treatment, the live birth rate per cycle showed significant positive correlation with sunshine hours during the month before the start of ovarian stimulation. The weather conditions or sunshine hours had no significant effect on spermatogenesis, which is consistent with our findings in the present study (Fig. 5A), although sperm mobility was not analyzed in the human study. Obviously, sperm is not the only factor driving the pregnancy outcome, our findings in the present study (Fig. 5B & C) suggest indoor lighting or deficiency in daylight exposure could have negative impact on male reproductive fitness.

### Conclusion and Translational Implications

More than a decade ago, Lucas et al. (2014) convened a transdisciplinary group of neuroscientists, vision researchers, lighting engineers, building scientists, and public health scientists to take what they called “the first important steps” toward measuring light in a way that accounts for what the human brain actually needs. The findings in the present study demonstrate that ecologically relevant lighting conditions differentially affect daily locomotor rhythms and physiological outcomes in a diurnal rodent model. Emulated daylight produced more robust daily synchronization than fluorescent light, a brief midday daylight pulse partially rescued daily rhythm amplitude, and hyperglycemia clustered among animals housed under fluorescent light. These findings provide controlled experimental evidence supporting the hypothesis that the indoor lighting environments in which most modern humans spend their waking hours are suboptimal for circadian function and, potentially, for metabolic and reproductive health. This study was conceived as use-inspired basic research (Stokes, 1997), motivated by both fundamental scientific questions and the practical needs of downstream stakeholders. The choice of CIE-standardized illuminants, the ecologically relevant condition hierarchy, and the inclusion of physiological endpoints beyond locomotor activity all reflect a deliberate effort to produce findings that can travel the health science nexus: from basic chronobiology, through building science, to public health guidance.

## Acknowledgements

The grass rat breeding colony was supported by NIH grants RNS139668 and RNS141983.

## Author contributions

ABK and LY designed the study; ABK, KL-D, MB, NL, MD, and NK conducted the experiments and analyzed data; ABK, HT, LC and LY wrote the manuscript.

## Statements and declarations

Not applicable.

## Ethical considerations

The Michigan State University Institutional Animal Care and Use Committee approved the experimental procedures used in this study (approval no. PROTO202500407) on Dec 03, 2025.

## Figures

**Suppl Figure 1.**
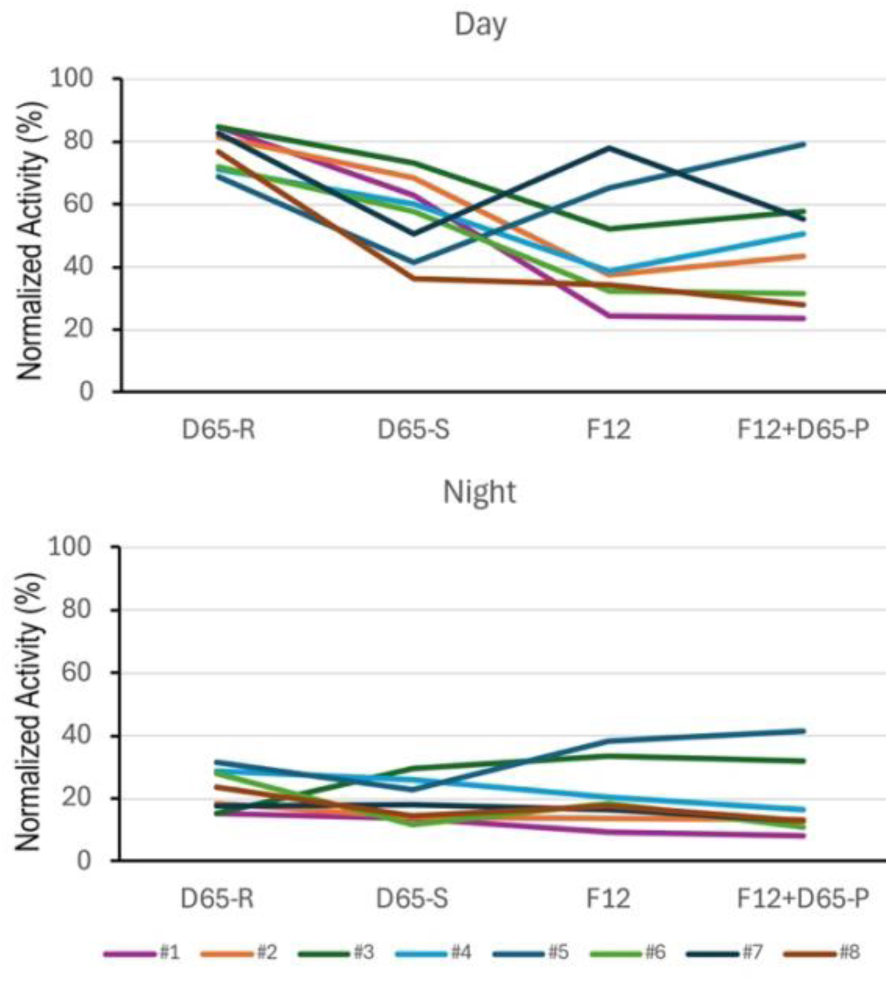
Day and night activity of each individual animals under the 4 lighting conditions. The activity in each phase and condition were normalized to total daily activities in D65-R.

**Suppl Figure 2.**
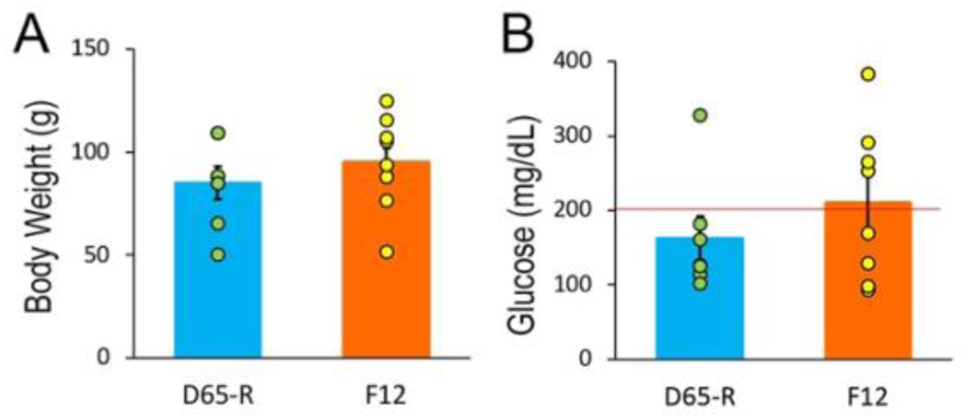
Body weight (A), and random blood glucose level (B) in animals housed under D65-R or F12 lighting conditions for 4 weeks. Glucose was measured from tail vein blood using a handheld glucometer while animals were briefly anesthetized with 5% isoflurane. Data are presented as mean ± SEM (n = 7-8/condition), with individual animals shown as filled circles. Red horizontal line in panel B indicates a blood glucose level of 200 mg/dL.

## Notes

### Competing Interest Statement

The authors have declared no competing interest.

